# Whole brain mapping of glutamate in adult and old primates at 11.7T

**DOI:** 10.1101/2021.09.10.459778

**Authors:** Clément M. Garin, Nachiket A. Nadkarni, Jérémy Pépin, Julien Flament, Marc Dhenain

**Affiliations:** Université Paris-Saclay, CEA, CNRS, Laboratoire des Maladies Neurodégénératives, 18 Route du Panorama, F-92265 Fontenay-aux-Roses, France; Commissariat à l’Energie Atomique et aux Energies Alternatives (CEA), Direction de la Recherche Fondamentale (DRF), Institut François Jacob, MIRCen, 18 Route du Panorama, F-92265 Fontenay-aux-Roses, France

**Author notes:** Correspondence Marc Dhenain, MIRCen, UMR CEA-CNRS 9199, 18 Route du Panorama, 92 265 Fontenay-aux-Roses CEDEX, France. These two authors participated equally to the work (Julien Flament, Marc Dhenain).

**Keywords:** Aging, Cerebral network, GluCEST, Glutamate, mouse lemur

## Abstract

Glutamate is the amino acid with the highest cerebral concentration. It plays a central role in brain metabolism. It is also the principal excitatory neurotransmitter in the brain and is involved in multiple cognitive functions. Alterations of the glutamatergic system may contribute to the pathophysiology of many neurological disorders. For example, changes of glutamate availability are reported in rodents and humans during Alzheimer’s and Huntington’s diseases, epilepsy as well as during aging.

Most studies evaluating cerebral glutamate have used invasive or spectroscopy approaches focusing on specific brain areas. Chemical Exchange Saturation Transfer imaging of glutamate (gluCEST) is a recently developed imaging technique that can map glutamate distribution in the entire brain with higher sensitivity and at higher resolution than previous techniques. It thus has strong potential clinical applications to assess glutamate changes in the brain. High field is a key condition to perform gluCEST images with a meaningful signal to noise ratio. Thus, even if some studies started to evaluate gluCEST in humans, most studies focused on rodent models that can be imaged at high magnetic field.

In particular, systematic characterization of gluCEST contrast distribution throughout the whole brain has never been performed in humans or non-human primates. Here, we characterized for the first time the distribution of the glutamate index in the whole brain and in large-scale networks of mouse lemur primates at 11.7 Tesla. Because of its small size, this primate can be imaged in high magnetic field systems. It is widely studied as a model of cerebral aging or Alzheimer’s disease. We observed high gluCEST contrast in cerebral regions such as the nucleus accumbens, septum, basal forebrain, cortical areas 24 and 25. Age-related alterations of this biomarker were detected in the nucleus accumbens, septum, basal forebrain, globus pallidus, hypophysis, cortical areas 24, 21, 6 and in olfactory bulbs. An age-related gluCEST contrast decrease was also detected in specific neuronal networks, such as fronto-temporal and evaluative limbic networks. These results outline regional differences of gluCEST contrast and strengthen its potential to provide new biomarkers of cerebral function in primates.

**Graphical abstract:** 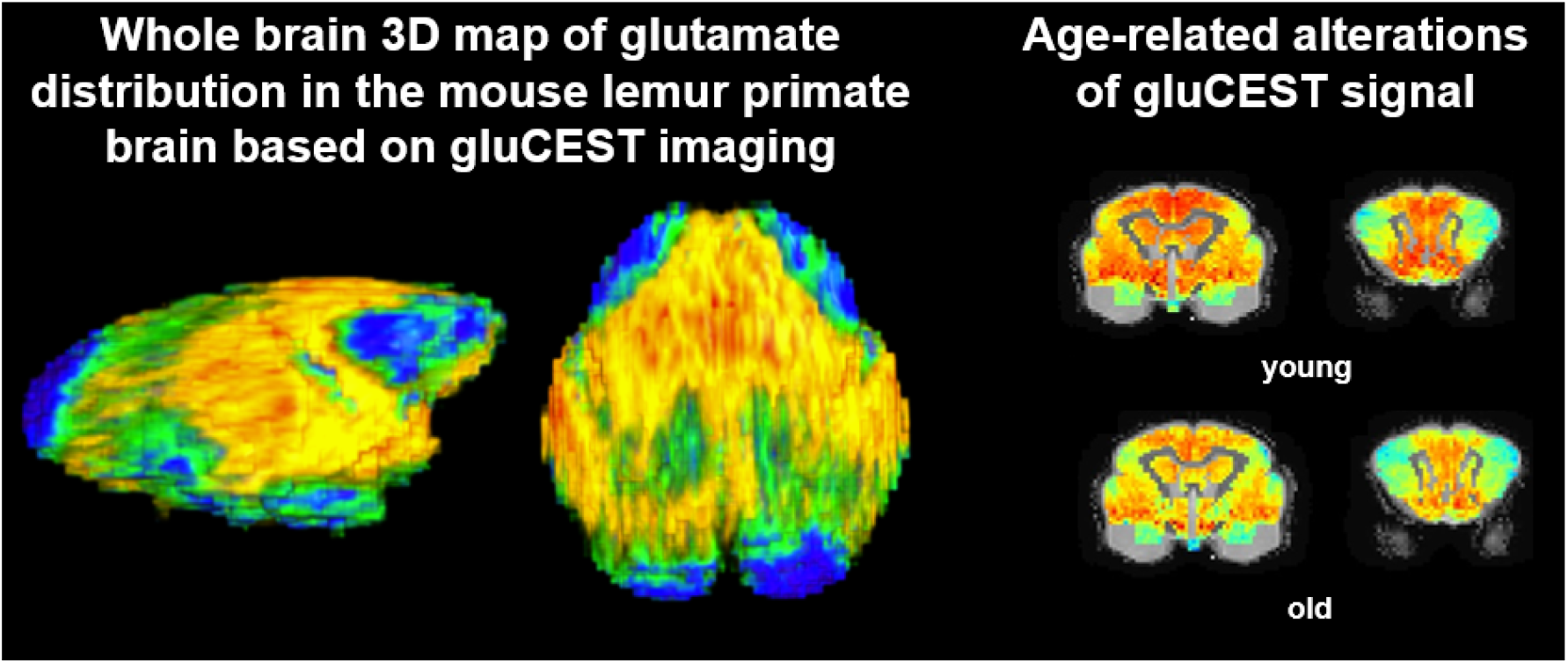

## 1. Introduction

Over the last century, human life expectancy has dramatically increased and the number of aged individuals is still rising. This trend results in the increased incidence of cerebral alterations that induce neurodegenerative diseases or mild cognitive/motor impairments which limit facets of daily living. This leads to a continuous demand for strengthening the research effort on mechanisms responsible for cerebral aging, and on therapies to prevent age-related diseases.

Glutamate is an essential amino acid of the brain metabolism and has the highest amino acid concentration of the brain (≈10 mmol/kg) (Greenamyre, 1986; Niciu et al., 2012). It is central to several metabolic pathways related to energy metabolism and oxidative stress (Sonnewald, 2014; Zhou and Danbolt, 2014). It is also the principal excitatory neurotransmitter in the brain and is involved in multiple cognitive functions. In normal conditions, glutamate is mostly located in the neurons (Cooper and Jeitner, 2016), neurotransmission being governed by few micromolar of extracellular glutamate.

Given, the importance of glutamate for the brain, *in vivo* markers for brain glutamate can have strong clinical application to provide better diagnostics and aid in the development of early therapeutic intervention. Many studies of glutamate concentrations were based on magnetic resonance spectroscopy. As examples, such studies highlighted age-related reduction of cerebral glutamate concentration in humans during aging (Roalf et al., 2020; Sailasuta et al., 2008) and these changes were associated with cognitive declines (Zahr et al., 2008). Changes of glutamate are also reported in neurodegenerative pathology as Alzheimer’s disease (Kirvell et al., 2006). One limitation of this method is that measurements are confined to relatively large voxels. Recent developments of Chemical Exchange Saturation Transfer of glutamate (gluCEST) imaging allow the mapping of glutamate distribution at the level of the whole brain (Cai et al., 2012; Carrillo-de Sauvage et al., 2015). CEST has been proposed to indirectly detect brain metabolites/macromolecules with labile protons (-OH, -NH2, -NH). Exchangeable protons of a metabolite (*e*.*g*., -NH2 amine group of glutamate) exhibit a resonance frequency that is shifted relative to bulk proton frequency. Likewise, exchangeable protons can be saturated using a frequency-selective radiofrequency pulse optimised to saturate amine group of glutamate, leading to a proportional decrease of water signal due to magnetization exchange (Roalf et al., 2017). Thus, the change in free water magnetization, while selectively saturating the exchangeable protons of the metabolite, represents an indirect measurement of the metabolite content and the specific glutamate contribution can be estimated by quantifying asymmetrical magnetization transfer ratio (Liu et al., 2010). Approximately 70% of GluCEST value has been shown to be due to glutamate, based on theoretical as well as experimental studies (Bagga et al., 2018; Cai et al., 2012).

High field is a key condition to perform gluCEST images with a meaningful signal to noise ratio. Thus, even if some studies started to evaluate gluCEST in humans (Nanga et al., 2018), most studies focused on rodent models that can be imaged at high magnetic field. Studies in these animals have shown that gluCEST can characterize glutamate alterations related to Alzheimer’s (Haris et al., 2013) or Huntington’s disease (Pepin et al., 2016) in mouse models. To our knowledge, systematic characterization of gluCEST contrast distribution have never been performed in humans or non-human primates. Here, we characterized for the first time the distribution of the glutamate index in the whole brain and in large-scale networks of mouse lemur primates (*Microcebus murinus*) at 11.7 Tesla.

This primate is characterized by its small size (typical length 12cm, 60-120g weight) and can thus easily be imaged with small bore high field MRI. As all non-human primates, it shares several genetic, physiological, and anatomical similarities with humans. It has a decade-long lifespan (Pifferi et al., 2018) and is widely used to study cerebral aging. As humans, it displays age-related cerebral atrophy that is associated with cognitive alterations (Picq et al., 2012; Sawiak et al., 2014), age-related accumulation of amyloid and tau lesions (Kraska et al., 2011), age-related increase of iron accumulation in the pallidum and substantia nigra (Dhenain et al., 1998). This animal is now used to characterize the impact of pathological processes as prediabetes on the brain (Djelti et al., 2016), to induce Alzheimer-like pathologies (Gary et al., 2019) as well as to assess interventions that can modulate cerebral aging (Pifferi et al., 2018).

First, we described variations of GluCEST contrast distribution in the whole brain. Second, we characterized brain regions associated with age-related reduction of gluCEST signal. Some functional regions are more intensely inter-connected between each other than to the rest of the brain, forming a level of functional organization called large-scale network. In the last part of the study, gluCEST contrast was evaluated within cerebral networks identified in mouse lemurs (Garin et al., 2021). The highest gluCEST signal was found in limbic networks and in high-level cortical networks such as fronto-temporal networks (FTN) and fronto-parietal network (FPN). FTN and FPN present homologies with human networks as the default mode network (DMN), the dorsal attentional network (DAN) and the executive control network (ECN) (Garin et al., 2021). Aging lowered gluCEST contrast in the FTN and evaluative limbic networks, which suggests that these two networks are prone to glutamate impairments.

## 2. Materials and methods

### 2.1. Animals and breeding

This study was carried out in accordance with the recommendations of the European Communities Council directive (2010/63/EU). The protocol was approved by the local ethics committees CEtEA-CEA DSV IdF (authorizations 201506051736524 VI (APAFIS#778)). All mouse lemurs studied were born in the laboratory breeding colony of the CNRS/MNHN in Brunoy, France (UMR 7179 CNRS/MNHN) and bred in our laboratory (Molecular Imaging Research Center, CEA, Fontenay-aux-Roses).

Thirty-three mouse lemurs (21 males and 12 females) were initially included in this study. Four animals that presented brain lesions or artefacted MR images were excluded from the analysis. Fourteen animals ranged from 1.3 to 3.8 years old (mean±SD: 2.1±0.8 years) were grouped together to form the “middle-aged lemurs cohort” (Table 1). Fifteen animals ranged from 8.0 to 10.8 years old (mean±SD: 8.8±1.1 years) were grouped together to form the “old lemurs cohort” (Table 1). Housing conditions were cages containing one or two lemurs with jumping and hiding enrichment, temperatures 24–26°C, relative humidity 55% and seasonal lighting (summer: 14 hours of light/10 hours of dark; winter: 10 hours of light/14 hours of dark). Food consisted of fresh apples and a homemade mixture of bananas, cereals, eggs and milk. Animals had free access to tap water. None of the animals had previously been involved in pharmacological trials or invasive studies.

**Table 1.**
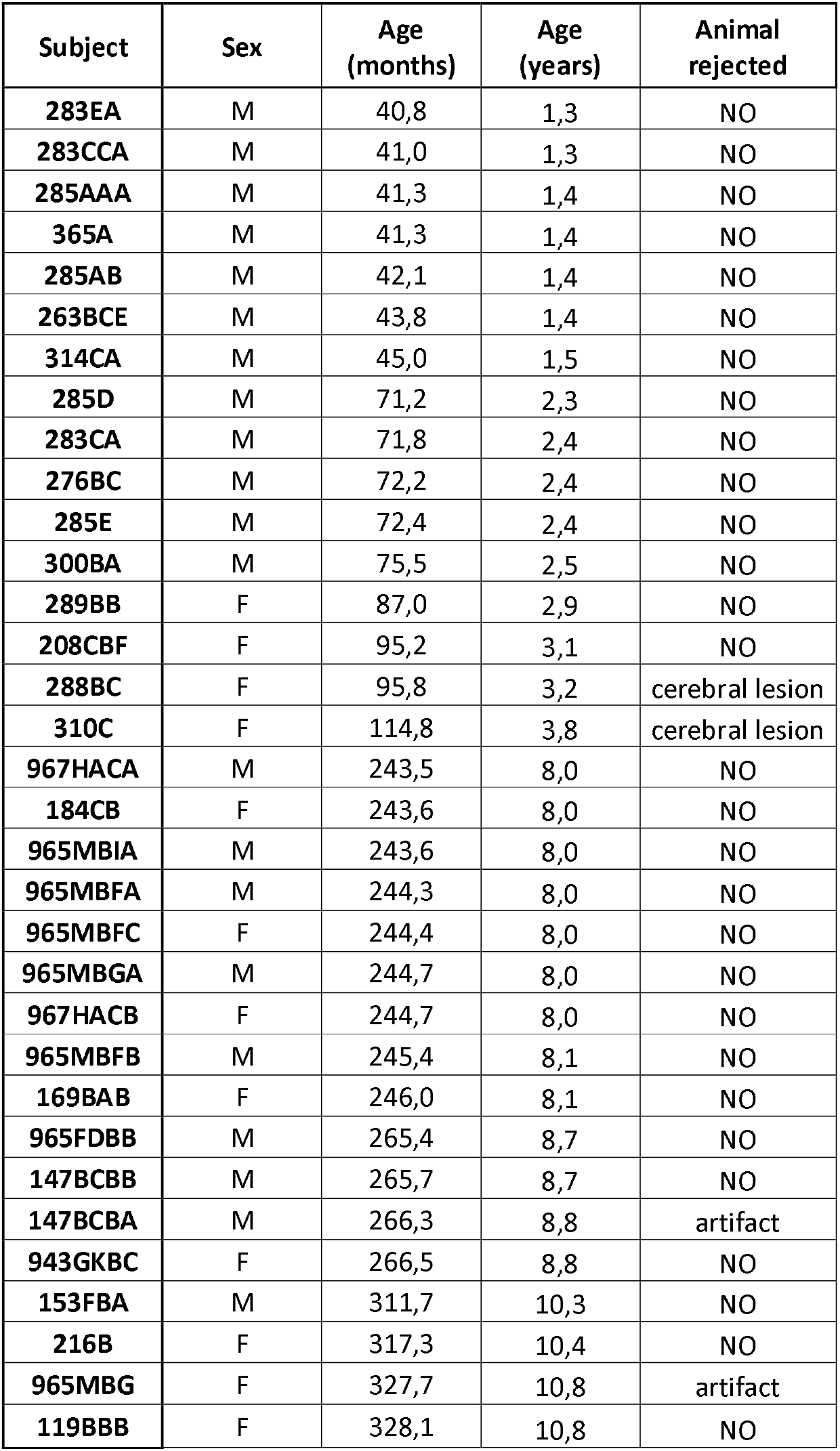
Cohort of mouse lemurs involved in the study.

| Subject | Sex | Age (months) | Age (years) | Animal rejected |
| --- | --- | --- | --- | --- |
| 283EA | M | 40,8 | 1,3 | NO |
| 283CCA | M | 41,0 | 1,3 | NO |
| 285AAA | M | 41,3 | 1,4 | NO |
| 365A | M | 41,3 | 1,4 | NO |
| 285AB | M | 42,1 | 1,4 | NO |
| 263BCE | M | 43,8 | 1,4 | NO |
| 314CA | M | 45,0 | 1,5 | NO |
| 285D | M | 71,2 | 2,3 | NO |
| 283CA | M | 71,8 | 2,4 | NO |
| 276BC | M | 72,2 | 2,4 | NO |
| 285E | M | 72,4 | 2,4 | NO |
| 300BA | M | 75,5 | 2,5 | NO |
| 289BB | F | 87,0 | 2,9 | NO |
| 208CBF | F | 95,2 | 3,1 | NO |
| 288BC | F | 95,8 | 3,2 | cerebral lesion |
| 310C | F | 114,8 | 3,8 | cerebral lesion |
| 967HACA | M | 243,5 | 8,0 | NO |
| 184CB | F | 243,6 | 8,0 | NO |
| 965MBIA | M | 243,6 | 8,0 | NO |
| 965MBFA | M | 244,3 | 8,0 | NO |
| 965MBFC | F | 244,4 | 8,0 | NO |
| 965MBGA | M | 244,7 | 8,0 | NO |
| 967HACB | F | 244,7 | 8,0 | NO |
| 965MBFB | M | 245,4 | 8,1 | NO |
| 169BAB | F | 246,0 | 8,1 | NO |
| 965FDBB | M | 265,4 | 8,7 | NO |
| 147BCBB | M | 265,7 | 8,7 | NO |
| 147BCBA | M | 266,3 | 8,8 | artifact |
| 943GKBC | F | 266,5 | 8,8 | NO |
| 153FBA | M | 311,7 | 10,3 | NO |
| 216B | F | 317,3 | 10,4 | NO |
| 965MBG | F | 327,7 | 10,8 | artifact |
| 119BBB | F | 328,1 | 10,8 | NO |

### 2.2. Animal preparation and MRI acquisition

Each animal was scanned twice with an interval of 6 months. All scanning was under isoflurane anaesthesia at 1.25-1.5% in air, with respiratory rate monitored to confirm animal stability until the end of the experiment. Body temperature was maintained by an air heating system at 32°C, inducing a natural torpor in mouse lemurs (Aujard and Vasseur, 2001). This has the advantage of allowing a low anaesthesia level without reawakening.

The MRI system was an 11.7 T Bruker BioSpec (Bruker, Ettlingen, Germany) running ParaVision 6.0.1 with a volume coil for radiofrequency transmission and a quadrature surface coil for reception (Bruker, Ettlingen, Germany).

Anatomical images were acquired using a T_2_-weighted multi-slice multi-echo (MSME) sequence: repetition time (TR) = 5000 ms, echo time (TE) = 17.5 ms, field of view (FOV) = 32 × 32 mm, 75 slices of 0.2 mm thickness, 6 echoes, inter echo time (IES) = 5 ms, resolution = 200 µm isotropic, acquisition duration 10 min.

GluCEST images covering the brain from prefrontal cortex to the occipital cortex were acquired with a 2D fast spin-echo sequence: TR = 20000 ms, TE = 6 ms, FOV = 24 × 24 mm, 12 slices of 1.5 mm thickness, resolution = 0.250 × 0.250 µm^2^, acquisition duration 33 m. The MAPSHIM routine was applied in a voxel encompassing the slices of interest in order to reach a good shim on gluCEST images. gluCEST images were preceded by a frequency-selective continuous wave saturation pulse and acquired with a saturation pulse applied during T_sat_ = 1 s, composed by 10 broad pulses of 100 ms, with 20 μs inter-delay and an amplitude B_1_ = 5 μT. The frequency of the saturation pulse Δω was applied in a range from −5 ppm to 5 ppm with a step of 1 ppm. In vivo, CEST contrast can be hampered by several competing factors such as direct saturation transfer (DS) of free water and background magnetization transfer (MT). Although we supposed DS symmetrical with respect to water frequency and suppressed by asymmetrical analysis its contribution to CEST contrast (Sun et al., 2005; van Zijl and Yadav, 2011; Zhou and van Zijl, 2006). B_0_ inhomogeneities were corrected using the Water Saturation Shift Reference (WASSR) method (Kim et al., 2009) with a saturation pulse applied around the water frequency (Δω in a range from -1 ppm to 1 ppm with a step of 0.1 ppm) with an amplitude B_1_ = 0.2 µT.

### 2.3. MRI pre-processing

CEST images were first processed pixel-by-pixel and analyzed using in-house programs developed on MATLAB software (MathWorks Inc., Natick, MA) used to generate Z-spectra by plotting the longitudinal magnetization as a function of saturation frequency. The specific glutamate contribution was isolated using Asymmetrical Magnetization Transfer Ratio (MTRasym) (Liu et al., 2010) and was calculated as follows: MTRasym(Δω) = 100 × (Msat(−Δω) − Msat(+Δω)) / Msat(−5 ppm), Msat(±Δω) being the magnetization acquired with saturation pulse applied at ‘+’ or ‘−’ Δω ppm. GluCEST images were calculated with Δω centered at ± 3 ppm. GluCEST image was converted into NIfTI-1 format.

Anatomical images were exported as DICOM files then converted into NIfTI-1 format. Then spatial pre-processing was performed using the python module sammba-mri (SmAll MaMmals BrAin MRI; http://sammba-mri.github.io) (Celestine et al., 2020) which, using nipype for pipelining (Gorgolewski et al., 2011), leverages AFNI (Cox, 1996) for most steps and RATS (Oguz et al., 2014) for brain extraction. Anatomical images were mutually registered to create a study template, which was further registered to a high-resolution anatomical mouse lemur template of the functional atlas (Garin et al., 2021). Then, gluCEST image were brought into the same space of the mouse lemur template by successive application of the individual anatomical to study template and study template to mouse lemur atlas transforms.

### 2.4. GluCEST contrast extraction and statistical analysis

To regional distribution of glutamate and differences of MTRasym between groups, we extracted gluCEST signal using a mouse lemur digital atlas (https://www.nitrc.org/projects/mouselemuratlas; (Nadkarni et al., 2019) as well as large scale networks maps (http://www.nitrc.org/projects/prim_func_2020/; (Garin et al., 2021) by applying NiftiLabelsMasker from Nilearn (Abraham et al., 2014)

Inter-individual differences between middle-aged and old groups were first evaluated using voxel wise statistical analysis (AFNI 3dttest++ (Cox, 1996)). Voxel wise analysis was performed only in regions with high gluCEST contrast (ten percent of the highest gluCEST voxels were kept). Cluster size (40 voxels) was estimated using AFNI (-Clusterize) with p<0.03 (uncorrected)). Using map_threshold from nilearn (Abraham et al., 2014), we extracted on the Ttest map cluster of voxels superior to 40 associated to p<0.03.

Then, we evaluated differences of gluCEST signal between middle-aged and old groups at different levels of segmentation (anatomy/network). Statistical differences between middle-aged and old animals were estimated using Kruskal-Wallis H-test. These complementary analyses allow to test the reproducibility of the voxel wise results as well as to evaluate the fit between anatomy/network and dysfunction.

### 2.5. Data availability

The template and atlas used in this study maps are available for download in NIfTI-1 format at https://www.nitrc.org/projects/mouselemuratlas. The code developed to create and manipulate the template has been refined into general procedures for registering small mammal brain MR images, available within a python module sammba-mri (SmAll-maMMals BrAin MRI; https://sammba-mri.github.io). MR images used in this study can be made available upon request after signing a data sharing agreement required by authors’ institution.

## 3. Results

### 3.1. GluCEST contrast distribution in middle-aged adult mouse lemurs

GluCEST images were acquired from 29 anaesthetised mouse lemurs (isoflurane 1.25-1.5%) at 11.7 Tesla (Table 1). First, GluCEST contrast was analysed in middle-aged adults (n=14, 1.3 to 3.8 year-old, mean±SD: 2.1±0.8 years, Table 1). Regions with the highest gluCEST contrast (average asymmetrical magnetization transfer ratio (MTRasym in %)) were found in subcortical regions and cortical regions (Fig. 1A, Fig. 2A). Subcortical regions with the highest contrast (median>16) were the nucleus accumbens, septum, basal forebrain, putamen, claustrum and caudate nucleus. Cortical regions with the highest contrast were frontal regions from the anterior cingulate areas (area 25 (subgenual area) and 24 (anterior cingulate cortex)) and area 4 (primary motor cortex). Cortical distribution of gluCEST contrast is further displayed on 3D views of the brain in Fig. 1B. This figure reflects average surface map of MTRasym, normalized by the average signal of the brain. This normalization allows visualizing variation strictly due to the distribution of gluCEST signal and not to inter-individual variations.

**Figure 1.**
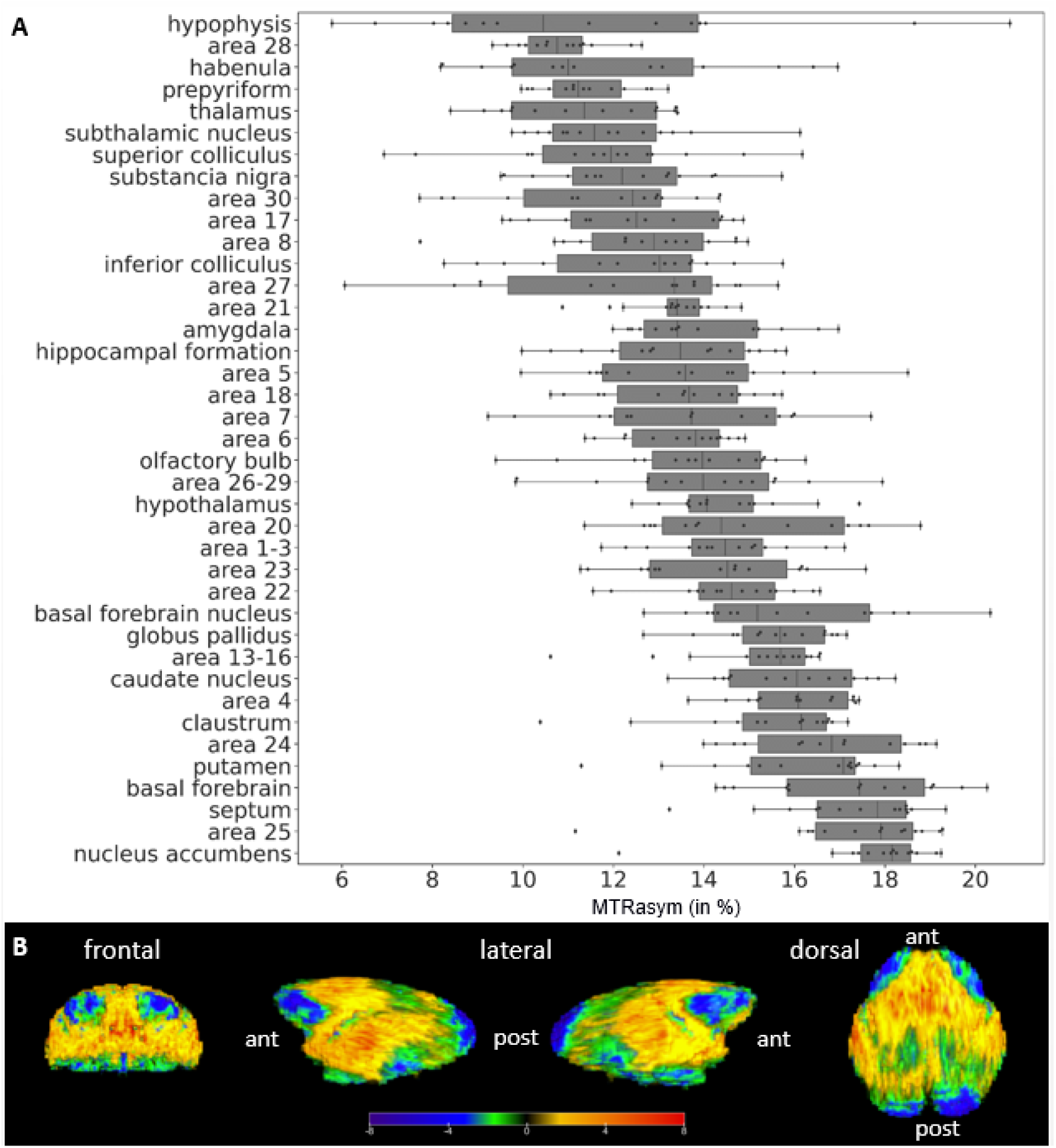
Regional MTRasym (in %) in middle-aged adult mouse lemurs. (A) MTRasym in different brain regions of middle-aged adult mouse lemurs (n=14) ranked based on their group median value. Elevated MTRasym is observed within regions encompassing the nucleus accumbens, septum, basal forebrain, putamen, and anterior cingulate regions (areas 24 and 25). (B) 3D surface map of the gluCEST signal showing regional signal variations. Color map: average surface map of MTRasym, normalized by the average signal of the brain ant: anterior part of the brain, post: posterior part of the brain.

**Figure 2.**
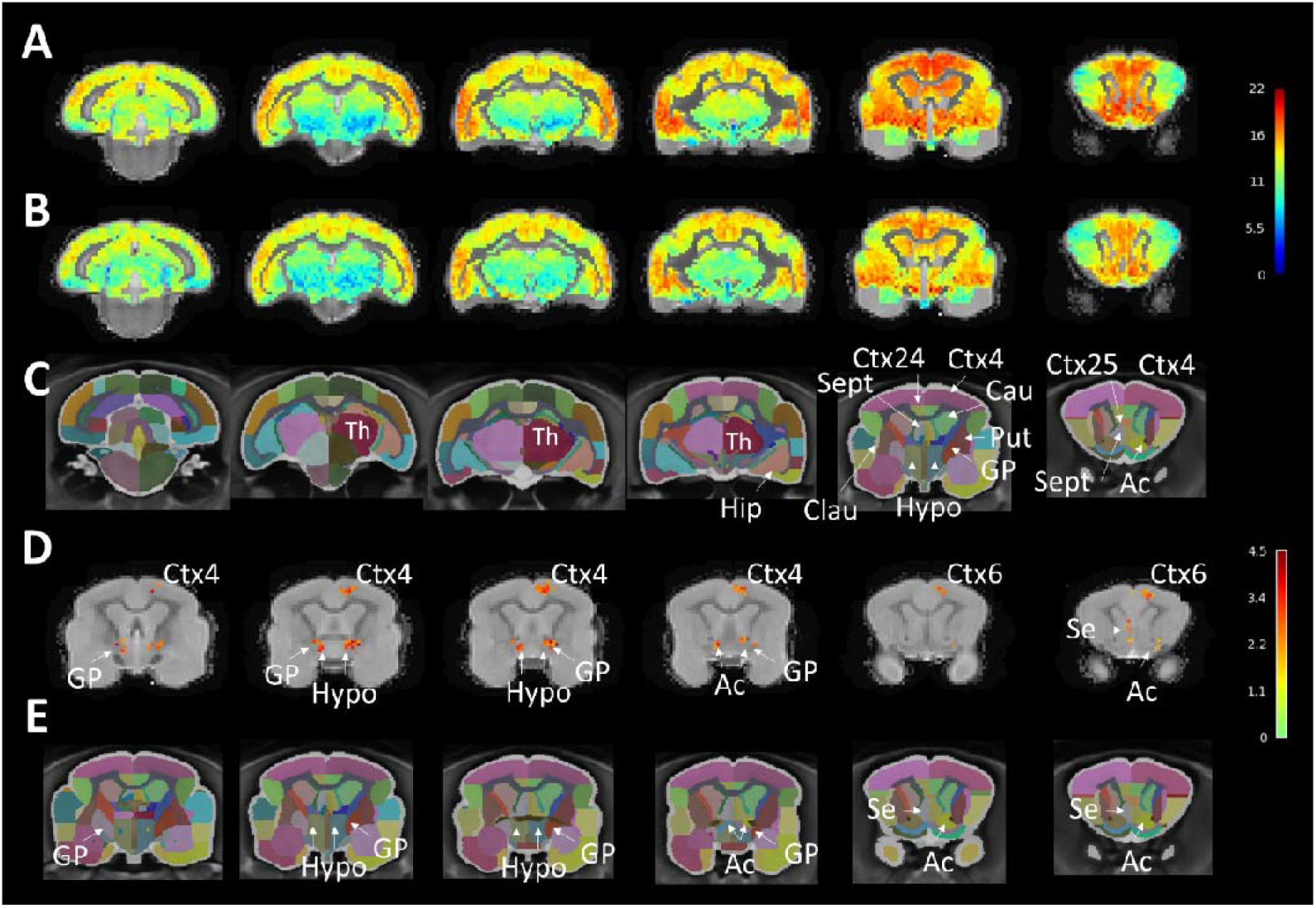
gluCEST contrast (MTRasym) in middle-aged and old mouse lemurs. Average MTRasym in middle-aged (A) and old mouse lemurs (B). Anatomical view of the mouse lemur brain corresponding to the displayed sections are displayed in C (Nadkarni et al., 2019). Significant statistical decrease of the gluCEST signal was detected using voxelwise analysis in the globus pallidus, nucleus accumbens, and to a lower extent hypothalamus and septum (D). Within the cortex, signal was reduced in area 4 (primary motor cortex) and area 6 (premotor cortex and supplementary motor area). The slices posterior and anterior to those shown in figure D and E did not display any significant differences. E displayed anatomical view of the mouse lemur sections shown in D (Nadkarni et al., 2019). Ac: Nucleus accumbens; Se: Septum; bf: Basal forebrain; Cau: Caudate nucleus; Clau: Claustrum; GP: Globus pallidus; Hip: Hippocampus; Hypo: Hypothalamus; Sept: Septum; Th: Thalamus. Scale bars in A-C: gluCEST contrast, in D: t-value.

GluCEST contrast was especially low in the entorhinal cortex (area 28), retrosplenial region (area 30), occipital cortex (area 17), thalamus and the colliculi.

### 3.2. Age-related regional alterations of GluCEST contrast

#### 3.2.1. Voxel based analysis

Inter-individual differences between maps of MTRasym in middle-aged (Fig. 2A) and old groups (Fig. 2B) were evaluated using voxel wise statistical analysis (Fig. 2D). This analysis highlighted a significant statistical decrease in voxels belonging to the globus pallidus, nucleus accumbens, and to a lower extent hypothalamus and septum. Within the cortex, signal was reduced in area 4 (primary motor cortex) and area 6 (premotor cortex and supplementary motor area) (Nadkarni et al., 2019) (Fig. 2D).

#### 3.2.2. Atlas based analysis

Comparison of gluCEST signal in middle-aged and old animals could also be performed using an atlas-based approach. This analysis confirmed that aging is associated with a decrease of glutamate level in the globus pallidus (*p = 0*.*00025*, Kruskal-Wallis H-test), nucleus accumbens (*p = 0*.*0013*, Kruskal-Wallis H-test). It detected additional subcortical regions with an age-related glutamate decrease as the hypophysis (*p = 0*.*0052*, Kruskal-Wallis H-test), septum (*p = 0*.*016*, Kruskal-Wallis H-test), and basal forebrain (*p = 0*.*032*, Kruskal-Wallis H-test). The hypothalamus that displayed focal reduction of gluCEST contrast was not detected by atlas-based approach. The cortical area 6 was detected with an atlas-based approach (*p = 0*.*029*, Kruskal-Wallis H-test) as well as additional cortical regions (*e*.*g*. area 25 (*p = 0*.*026*, Kruskal-Wallis H-test), area 21 (middle temporal area, *p = 0*.*0040*, Kruskal-Wallis H-test)) (Fig. 3).

**Figure 3.**
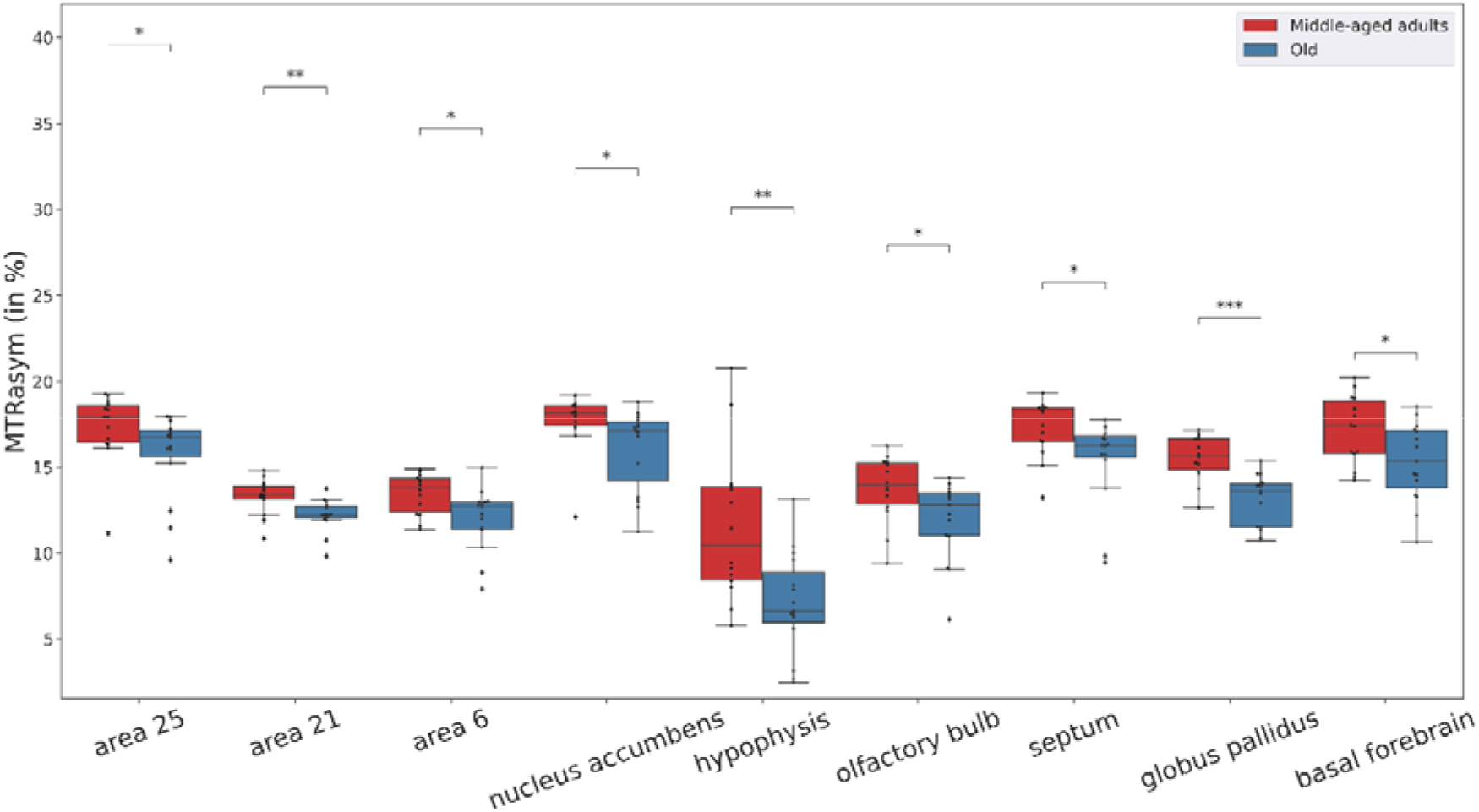
Significant differences of gluCEST signal between middle-aged and old mouse lemurs as detected by atlas-based approach. *: p <0.05; **: p <0.01, ***: p < 0.001, ****: p < 0.0001.

#### 3.2.3. Age-related impairments of gluCEST contrast involves large scale networks

We then evaluated the extracted contrast within cerebral networks previously reported in mouse lemurs (Garin et al., 2021). GluCEST differences between middle-aged and old animals were found in the fronto-temporal network (Fig. 4; *p = 0*.*016*, Kruskal-Wallis H-test) and the evaluative limbic network (*p = 0*.*026*, Kruskal-Wallis H-test). The fronto-temporal network involved the frontal anterior medial and lateral regions, the precentral cortex, all the temporal regions, the parietal posterior cortex, the anterior and medial cingulum cortices, and the insular cortex. The evaluative-limbic network embedded limbic structures, the insula, as well as subcortical structures.

**Figure 4.**
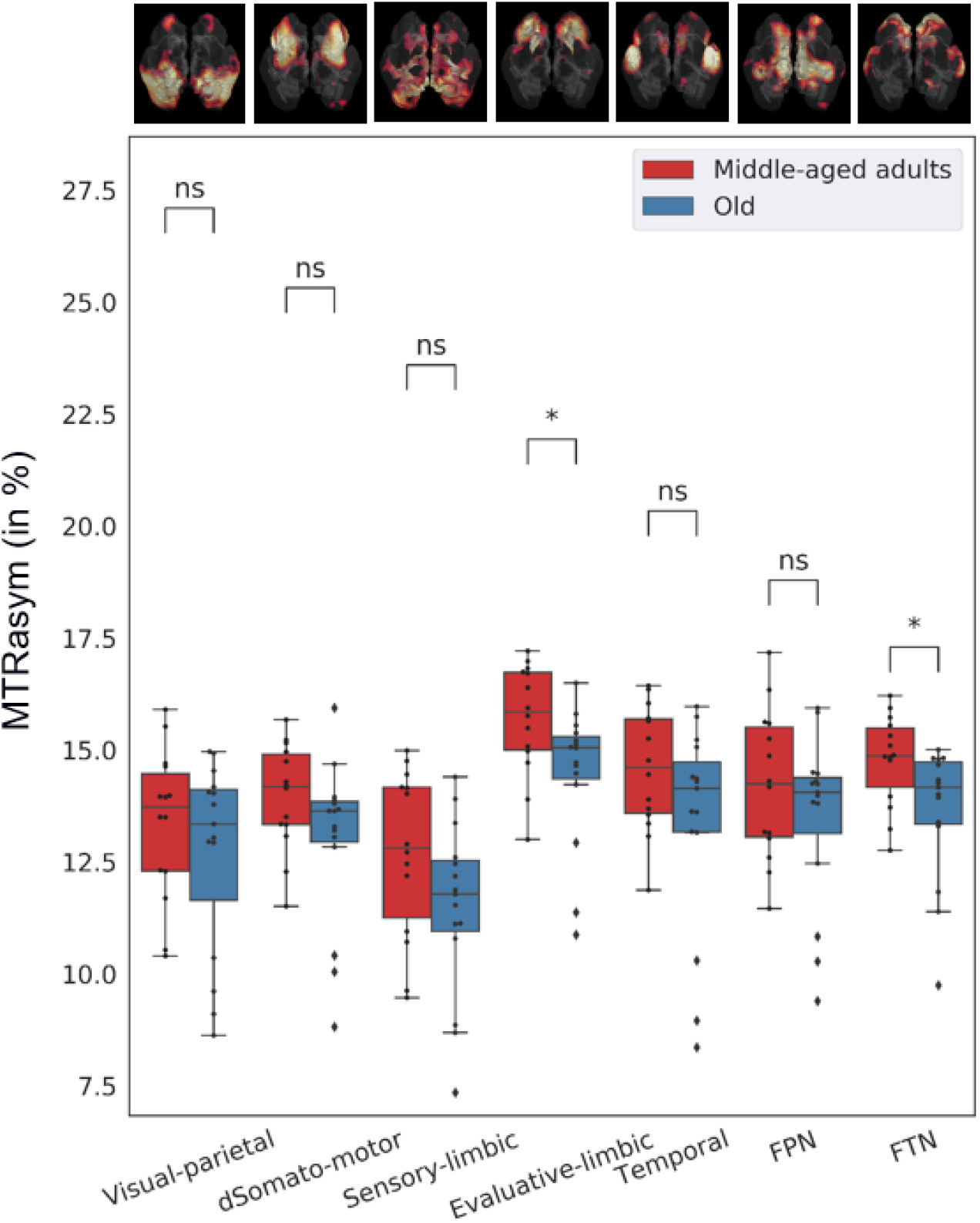
gluCEST signal within large-scale network of middle-aged and old mouse lemurs. Signal was lower in the evaluative limbic network (*p = 0*.*026*, Kruskal-Wallis H-test) and in the fronto-temporal network (FTN, *p = 0*.*016*, Kruskal-Wallis H-test) of old animals when compared to middle-aged animals. At the top of the figure, large-scale networks are illustrated using images based on Garin et al. (Garin et al., 2021). *: p < 0.05; **: p < 0.01, ***: p < 0.001, ****: p < 0.0001. FPN: Fronto-parietal network.

## 4. Discussion

This study characterized for the first-time glutamate distribution and its age-related changes in the whole brain of a non-human primate. High field is a key condition to perform gluCEST images with a meaningful signal to noise ratio. Given its small body size, the mouse lemur could fit in small-bore high field MRI scanner, which opened the possibility for gluCEST imaging in a non-human primate.

### 4.1. Distribution of gluCEST contrast in normal middle-aged animals

Glutamate is an essential amino acid and has the highest amino acid concentration of the brain. It is involved in several metabolic pathways related to energy metabolism and oxidative stress (Sonnewald, 2014; Zhou and Danbolt, 2014). It is also the major excitatory neurotransmitter in the central nervous system (Cooper and Jeitner, 2016).

The main glutamate pathways involve projections **i**. from the prefrontal cortex to the striatum (caudate nucleus and putamen: cortico-striatal pathway) and the nucleus accumbens (cortico-accumbens pathway), **ii**. from the prefrontal cortex to brain-stem nuclei (cortico-brainstem pathway), **iii**. from the prefrontal cortex to the thalamus (cortico-thalamic pathway). **iv**. from the thalamus to the cortex (thalamo-cortical pathway), and **v**. from the cortex to the cortex (cortico-cortical pathways) (Greenamyre, 2001; Schwartz et al., 2012).

We found high glutamate levels in several regions involved in these pathways. In cortical regions, high gluCEST contrast was observed in the anterior cingulate regions (Brodmann area 25 (subgenual area) and 24 (anterior cingulate cortex)) and premotor cortex (area 6). These regions fit with human “cortico-striatal/accumbens/thalamic/brain stem” pathways. Most of the subcortical regions in which we detected high gluCEST contrast were also involved in these pathways. Indeed, high levels were found in the putamen and caudate nucleus, involved in the cortico-striatal pathway. Numerous studies outlined the strong cortical glutamatergic projections on these structures (Galvan et al., 2006; Lanciego et al., 2012). The nucleus accumbens involved in the cortico-striatal pathway had also high glutamate levels. Some other regions, not as involved in these pathways as the septum, basal forebrain, however also had high gluCEST contrasts. Of note, neurotransmission is governed by few micromolar of extracellular glutamate so one can hypothesize that the transmitter pool of glutamate contributes very slightly to the gluCEST contrast. Thus, even if gluCEST changes occur close to glutamate pathways, they do not directly reflect changes of neurotransmitters, but mainly pools of glutamate involved in metabolic pathways.

### 4.2. Age-related changes of gluCEST contrast

In the second part of our study, we showed that aging leads to reduction of gluCEST contrast in subcortical regions as the nucleus accumbens, septum, basal forebrain, globus pallidus, hypophysis and to a lower extent hypothalamus. In the cortex, the reduction of gluCEST contrast occurred in frontal regions (anterior cingulate regions (areas 24-25), area 6), temporal regions (area 21). Parietal regions (area 4) also displayed some age-related changes. Minor differences between voxel-based and atlas-based analysis were observed. They can easily be explained by i. the exclusion of low gluCEST contrast areas in voxel-based analysis ii. the divergence of resolution (voxel vs anatomical region) iii. Atlas-based analysis results are displayed without correction for multiple comparisons. The few differences between the two analyses do not modify the interpretation of our study but their similarities strengthen the robustness of the detected alterations. In humans, an age-related glutamate reduction has been described in similar regions as in mouse lemurs. In particular, a reduction of glutamate was highlighted in the globus pallidus and the putamen using MR spectroscopy based on voxels that embedded these regions. Lower glutamate levels in these regions were associated to cognitive decline (Zahr et al., 2008). An age-related reduction of glutamate levels were also reported in some of the motor cortex in humans (Kaiser et al., 2005).

In the last part of the study, we showed that age-related changes of gluCEST contrast involved two large-scale networks: the fronto-temporal network and the evaluative limbic network. The fronto-temporal network is partly homologous to the human executive control network, DMN, salience and fronto-temporal networks (Garin et al., 2021). It was suggested that this network is involved in interoception and in self-referential decisions (Garin et al., 2021). We can speculate that age-related glutamate in this network may impair these functions, although it is difficult to test them in lemurs. The evaluative limbic network subserves behavioural responses to the positive/negative valence of stimuli as well as to the internal state of the body (Garin et al., 2021). In humans, several reports highlighted age-related impairments of large-scale networks (Ferreira and Busatto, 2013; Zhang et al., 2014). Impairments of white matter and dopamine as well as accumulation of amyloid in the brain have been suggested as potential culprits for these changes (Ferreira and Busatto, 2013). Our study suggests, that for at least some networks, glutamate level changes can also be involved in network impairments, although the impact of glutamate changes on network function will have to be further investigated in the future.

## 5. Conclusion

This study, based on gluCEST MRI, characterized glutamate at the whole brain level in a primate and reported age-related cerebral alterations in several key regions. This technic appears to be an essential tool to better understand the various mechanisms altering the brains and promises multiple applications for the diagnosis, follow up, treatment efficiency of neurodegenerative diseases.

## 6. Acknowledgements

We thank the France-Alzheimer Association, Plan Alzheimer Foundation and the French Public Investment Bank’s “ROMANE” program for funding this study. The 11.7T MRI scanner was funded by a grant from NeurATRIS: A Translational Research Infrastructure for Biotherapies in Neurosciences (“Investissements d’Avenir”, ANR-11-INBS-0011). C.G. was financed by the French Ministère de l’Enseignement Supérieur, de la Recherche et de l’Innovation.

## 8. Author contributions

C.M.G., J.F., and M.D. contributed to the study conception and design. J.P. and J.F. designed the gluCEST sequences, C.M.G. and N.A.N. designed registrations strategies and pipelines. C.M.G., J.F., and M.D. wrote the manuscript. All authors commented on previous versions of the manuscript. All authors read and approved the final manuscript.

